# TemBERTure: Advancing protein thermostability prediction with Deep Learning and attention mechanisms

**DOI:** 10.1101/2024.03.28.587204

**Authors:** Chiara Rodella, Symela Lazaridi, Thomas Lemmin

**Author notes:** Authors contributed equally (Chiara Rodella, Symela Lazaridi).

## Abstract

Understanding protein thermostability is essential for various biotechnological and biological applications. However, traditional experimental methods for assessing this property are time-consuming, expensive, and error-prone. Recently, the application of Deep Learning techniques from Natural Language Processing (NLP) was extended to the field of biology, with an emphasis on protein modeling. From a linguistic perspective, the primary sequence of proteins can be viewed as a string of amino acids that follow a physicochemical grammar.

This study explores the potential of Deep Learning models trained on protein sequences to predict protein thermostability which provide improvements with respect to current approaches. We implemented TemBERTure, a Deep Learning framework to classify the thermal class (non-thermophilic or thermophilic) and predict and melting temperature of a protein, based on its primary sequence. Our findings highlight the critical role that data diversity plays on training robust models. Models trained on datasets with a wider range of sequences from various organisms exhibited superior performance compared to those with limited diversity. This emphasizes the need for a comprehensive data curation strategy that ensures a balanced representation of diverse species in the training data, to avoid the risk that the model focuses on recognizing the evolutionary lineage of the sequence rather than the intrinsic thermostability features. In order to gain more nuanced insights into protein thermostability, we propose leveraging attention scores within Deep Learning models to gain more nuanced insights into protein thermostability. We show that analyzing these scores alongside the 3D protein structure could offer a better understanding of the complex interplay between amino acid properties, their positioning, and the surrounding microenvironment, all crucial factors influencing protein thermostability.

This work sheds light on the limitations of current protein thermostability prediction methods and introduces new avenues for exploration. By emphasizing data diversity and utilizing refined attention scores, future research can pave the way for more accurate and informative methods for predicting protein thermostability.

**Availability and Implementation:** TemBERTure model and the data are available at https://github.com/ibmm-unibe-ch/TemBERTure

## 1. Introduction

Biocatalysts have become integral to numerous industrial processes, ranging from pharmaceutical production to food processing and biofuels production^1–3^. In these applications, protein thermostability plays a crucial role^4,5^. Proteins that endure high temperatures are essential for accelerating and enhancing chemical reactions, leading to reduced production costs^2^. However, exposure to elevated temperatures can cause denaturation and loss of biological activity^6^, underscoring the importance of improving our understanding of protein thermostability.

Despite notable progress in experimental techniques for measuring protein thermostability, the process remains time-consuming and challenging to scale up, resulting in limited data on protein thermostability^7^. Currently, ProThermDB is the largest dataset of experimental thermodynamic data for protein stability^8^, encompassing a comprehensive collection of 32,000 proteins, of which 38% are wild-type sequences and 51% single point mutations. In recent developments, novel experimental techniques have emerged that allow for the determination of the thermal stability of proteins across the entire genome of a cell. These techniques involve the integration of mass spectrometry with limited proteolysis^9^, or liquid chromatography^10^. In addition to experimental techniques, the growth temperature of organisms is commonly employed as a proxy for protein thermostability^11–15^.

By comparing statistical data from thermophilic and non-thermophilic protein sequences, key features associated with thermostability have been identified, including higher proportions of hydrophobic and charged residues, and specific dipeptide motifs of thermophilic proteins^13,16– 19^. A higher occurrence of hydrogen bonds, salt bridges, disulfide bonds, and hydrophobic interactions is also observed in thermophilic proteins^20–23^.

Extensive research has led to the development of several machine learning models aimed at predicting protein thermostability, treating it as a classification task^15,24–31^. Early models like Thermopred employed a Support Vector Machines (SVM) classifier trained on a dataset of 793 non-thermophilic and 915 thermophilic protein sequences^15^, which became a foundation for training subsequent models^29,30^. An expanded version of this dataset, consisting of 1368 thermophilic and 1443 non-thermophilic proteins, was utilized for training the iThermo model, a multi-layer perceptron (MLP)^12^ and the Sapphire framework, a staking-based ensemble model^31^. Other models have approached the problem as a regression task to directly predict the melting temperature^32,33^.

Transformer-based models such as Bidirectional Encoder Representations from Transformers (BERT)^34^, have improved Natural Language Processing (NLP). By considering proteins as a string of amino acids, NLP can be applied to biology and more specifically to protein modeling and classification. ProtTrans^35^, a family of models including protBERT, leverages transformers to extract protein characteristics from sequence data. BertThermo^36^ uses the protBERT embeddings with classical machine learning models for thermophilicity classification, whereas DeepSTABp incorporates ProtTrans-XL embeddings and growth temperature to predict protein melting temperature^37^. Similarly, TemStaPro^38^ is an ensemble of models incorporating ProtT5-XL^35^ embeddings to feed-forward densely connected neural network models, and ProLaTherm^39^ integrates the encoder part of a T5-3B ^40^ model with ProtT5-XL^35^ as the feature extractor.

To overcome the shortcomings of present model approaches, we developed TemBERTure, a deep-learning package for protein thermostability prediction. It consists of three components: (i) TemBERTure_DB_, a large curated database of thermophilic and non-thermophilic sequences, (ii) TemBERTure_CLS_, a classifier and (iii) TemBERTure_Tm_, a regression model, which predicts, respectively, the thermal class (non-thermophilic or thermophilic) and melting temperature of a protein, based on its primary sequence. Both models are built upon the existing protBERT-BFD language model^35^ and fine-tuned through an adapter-based approach^41,42^. Our findings demonstrate the remarkable capability of Deep Learning to differentiate protein classes based on their sequences. However, it also highlights the current limitations imposed by the currently available data. Despite these limitations, the insights gained from the attention scores within these models offer promising clues to unraveling the underlying mechanisms of protein thermostability. This has the potential to unlock new avenues for research in biotechnology and protein engineering.

## 2. Results

### 2.1 TemBERTure_DB_

To train our Deep Learning models for predicting protein thermostability, we curated TemBERTure_DB_, a comprehensive dataset built upon the Meltome Atlas^10^ that includes data for over 48,000 proteins across 13 different species (Figure 1A). We further enriched it with all protein sequences from UniProtKB for each organism^43^. This initially resulted in a highly imbalanced dataset with only 44,000 sequences from thermophilic organisms (growth temperature above 60°C) compared to 4.3 million sequences from non-thermophilic organisms. To address this imbalance, we incorporated thermophilic proteomes from BacDive, adding 0.9 million sequences^44^. However, the thermophilic dataset remained biased towards bacterial and archaeal sequences. Therefore, we included similar bacterial sequences (< 30°C growth) with high identity (>80%) to thermophiles. This added valuable non-thermophilic examples outside the target class, for a more challenging training set (Table S1).

**Figure 1.**
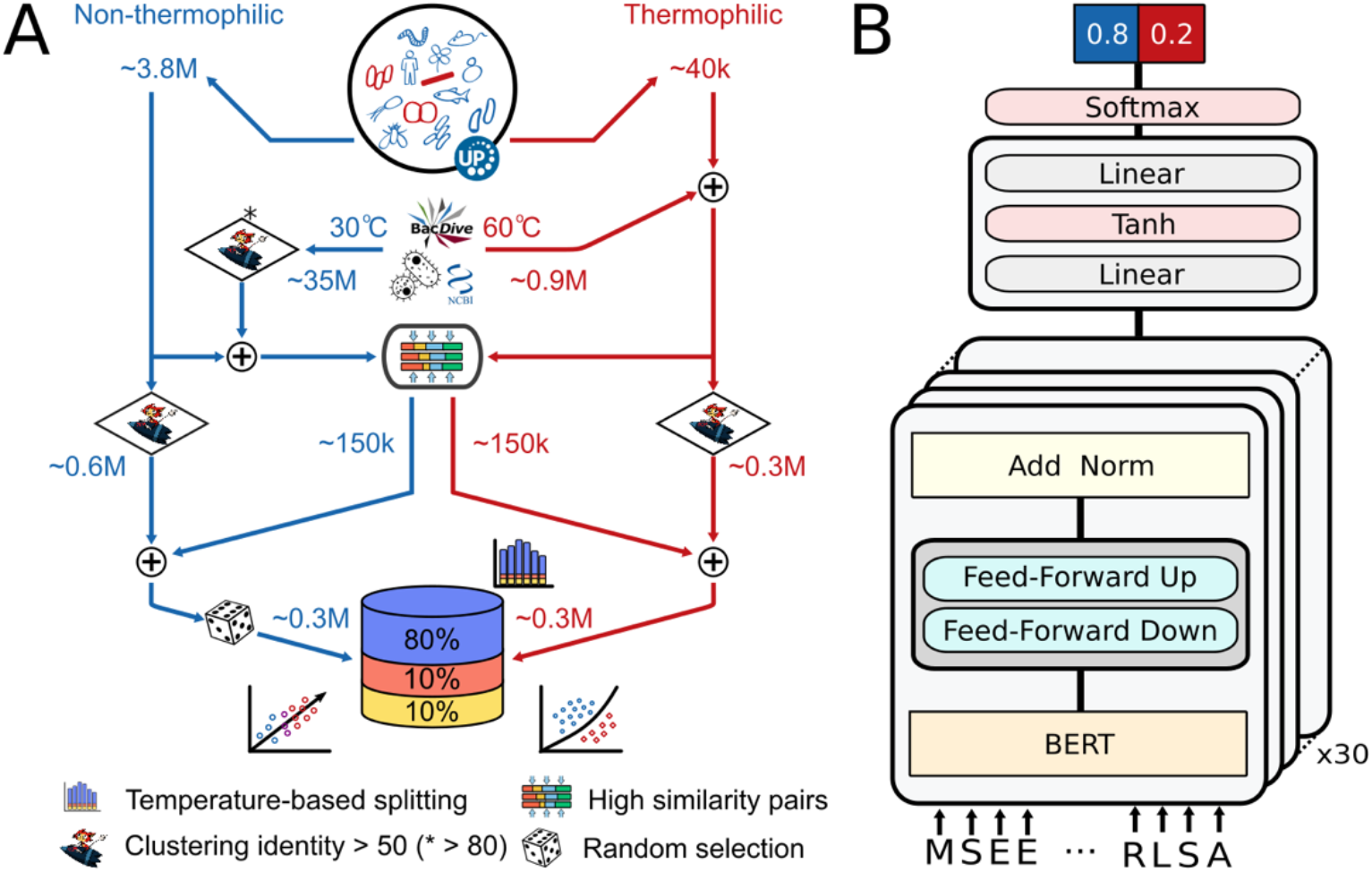
TemBERTure database creation and model architecture. (A) TemBERTureDB creation pipeline: Protein sequences from organisms within the Meltome Atlas were retrieved from the UniProt database and categorized based on their thermophilicity (red: thermophilic, blue: non-thermophilic). Additional sequences were then collected from BacDive and NCBI databases at various temperature thresholds to augment the dataset. The final database comprises approximately 0.3 million each for thermophilic and non-thermophilic proteins, further divided into training, testing, and validation sets that are representative of the temperature distribution. (B) TemBERTureCLS model architecture was based on the prot_bert_bfd framework, with lightweight bottleneck adapter layers inserted between each transformer layer (shown in gray). The model takes a protein sequence as input and outputs a score indicating the classification score of the sequence being thermophilic or non-thermophilic.

To ensure that both classes contained diverse protein families and folds, we clustered each class separately using MMseqs^45^, resulting in a balanced dataset of 300,000 sequences per class. We partitioned it into training, validation, and test sets at an 80:10:10 ratio, ensuring that sequences with high similarity remained within the same split, to avoid information leakage. To enhance model learning and generalization, pairs of highly similar sequences from different classes were exclusively reserved for training, effectively bridging the gap between thermophilic and non-thermophilic sequences (Table S2).

### 2.2 TemBERTure_CLS_

TemBERTure_DB_ served as the training dataset for TemBERTure_CLS_, a sequence-based classifier designed to predict the thermal class of a protein solely from its amino acid sequence (Figure 1B). TemBERTure_CLS_ leveraged protBERT-BFD, a pre-trained protein language model^35^, and utilized adapter layers^41,42^ for efficient task-specific learning. This approach offers faster (up to 50%) and more robust training (avoiding catastrophic forgetting) than full finetuning, thus enabling rapid model experimentation and optimization without sacrificing performance.

TemBERTure_CLS_ achieved an overall accuracy of 0.89, a F1-score of 0.9, and a Matthews Correlation Coefficient (MCC) of 0.78, with balanced predictive performance across both classes (0.88 and 0.90 F1-score for non-thermophilic and thermophilic sequence respectively). Low standard deviation across multiple trained models confirms robust training. We therefore chose to retain the initially trained model as the final TemBERTure_CLS_ model. When comparing the performance of TemBERTure_CLS_ to state-of-the-art models, we observed that many of the latter tend to overpredict the non-thermophilic class (Figure 2). Despite achieving a competitive average precision of 0.97 for thermophilic sequences, their recall fell below 0.7, resulting in numerous misclassifications of non-thermophilic proteins. This highlights the limitations in the generalizability of current methods (Table S3).

**Figure 2.**
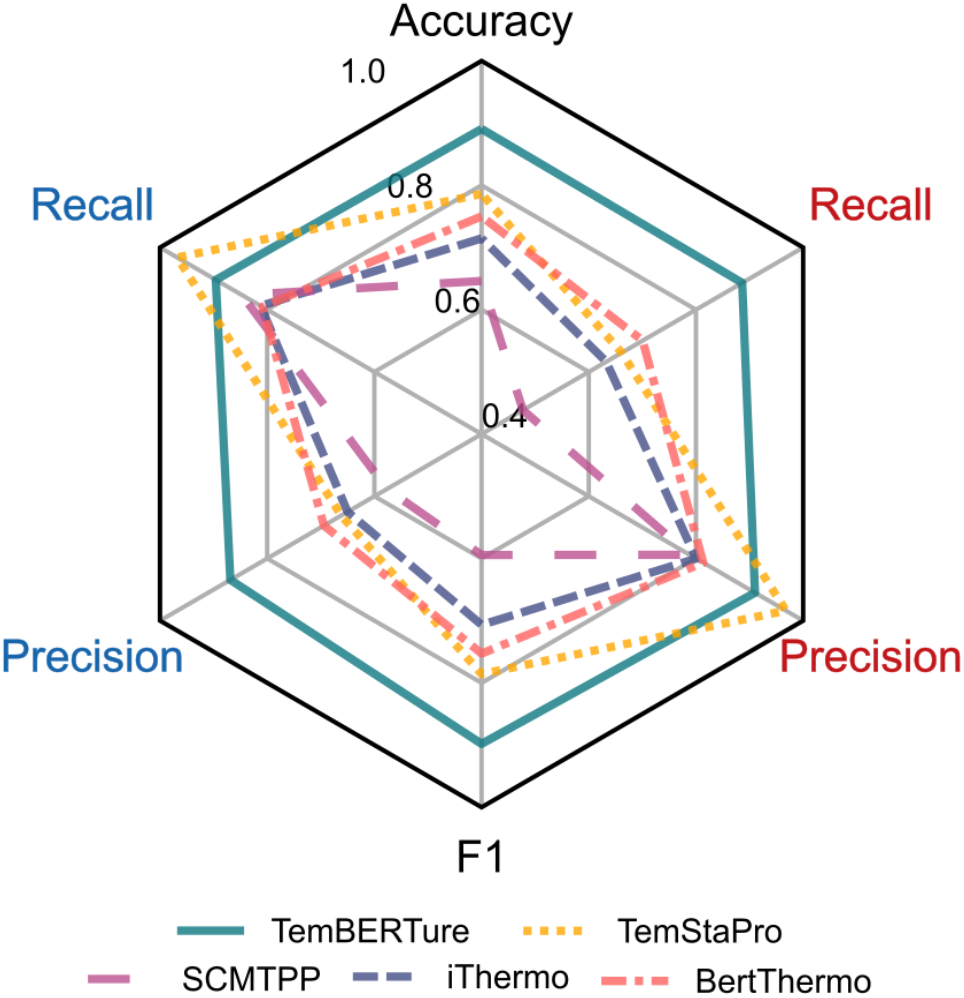
Comparison of TemBERTure_CLS_ with state-of-the art models on the TemBERTure_DB_ test set. Recall and Precision are shown separately for thermophilic (red) and non-thermophilic (blue) thermal categories.

To assess the generalization of TemBERTure_CLS_, we tested it on the widely used iThermo dataset^12^ and the TemStaPro test set^38^. After removing similar sequences (over 50% identity), the final test sets contained 65 and 1495 thermophilic sequences and 505 and 10849 non-thermophilic sequences for iThermo and TemStaPro, respectively. TemBERTure_CLS_ maintained high accuracy, achieving 86% on iThermo and 83% on TemStaPro (Table S4). To explore TemBERTure_CLS_ ability to perform on sequences from novel organisms, we created a new test set with sequences from organisms in the BacDive database^44^. Although non-thermophilic sequence precision remained high (0.81), precision for thermophilic sequences dropped (0.74), suggesting limitations in generalizing to completely new organisms.

To further investigate this observation, we trained separate models, with the same architecture as TemBERTure_CLS_, with two distinct datasets: one derived from BacDive^44^, focusing solely on bacterial and archaeal organisms, and another one from the Meltome Atlas^10^, augmented solely with thermophilic sequences (Tables S5 and S6). Each model performed well on the dataset derived from the same source as its training data. However, performances dropped significantly when tested on the other datasets (Figure 3). These variations were less pronounced for the thermophilic class, most likely because all datasets used BacDive for selecting thermophilic organisms. In contrast, the non-thermophilic class exhibited greater performance variations. The BacDrive-trained model’s performance dropped significantly, when tested on the TemBERTure_DB_ or Meltome_DB_ data (almost random classifications), whereas TemBERTure_CLS_ and the Meltome-trained model maintained comparable performance across all datasets, indicating the necessity of using diverse training datasets to improve generalizability.

**Figure 3.**
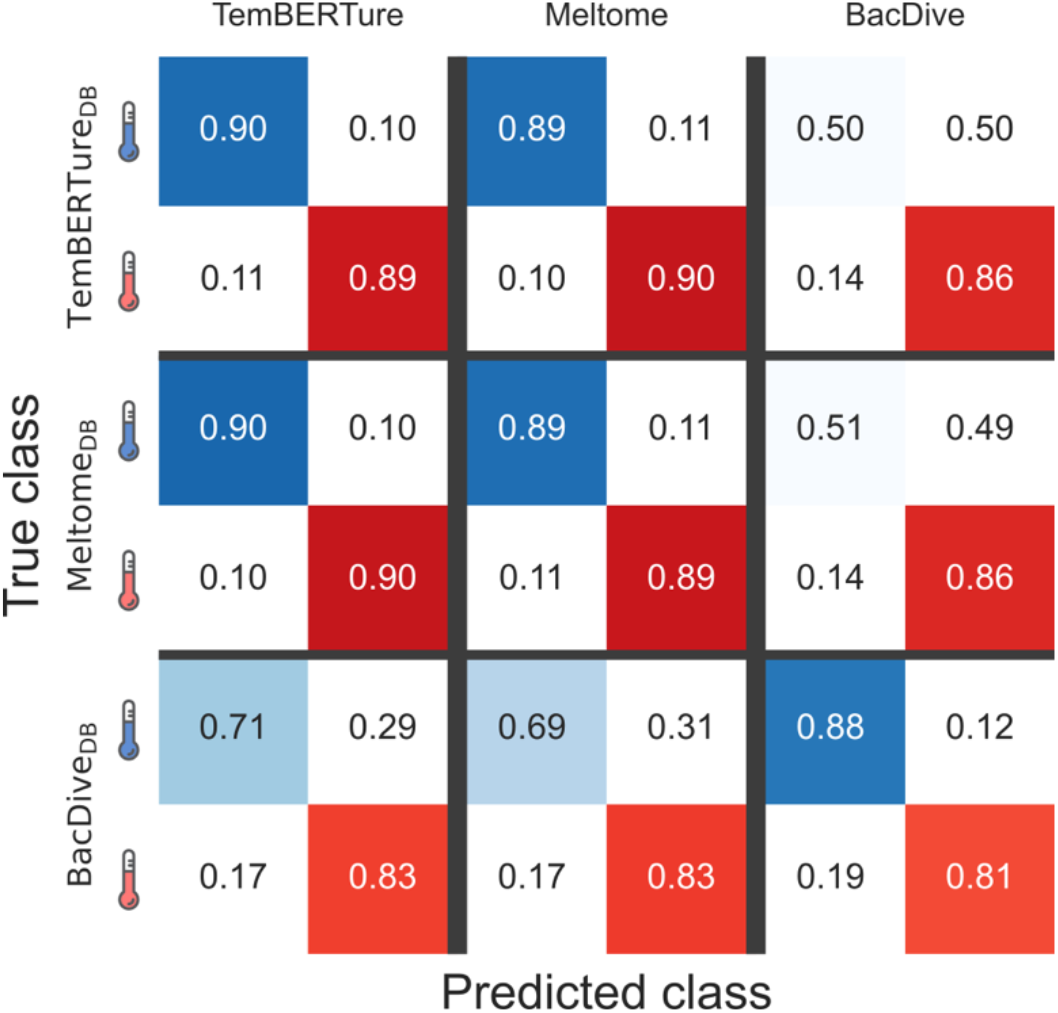
Impact of dataset curation on model performance. Confusion matrix comparing the performance of the TemBERTure_CLS_ model with models trained on data derived from only BacDive and Meltome. The evaluation is performed on three separate test sets: TemBERTure_DB_, BacDive_DB_ and Meltome_DB_ test sets. Each cell in the matrix represents the proportion of predictions made by a specific model on a specific test set. Shades of blue indicates correct predictions for the non-thermophilic category, while shades of red represents the performance for thermophilic sequences. Off-diagonal entries indicate instances of misclassification.

### 2.3 TemBERTure_Tm_

Building on these promising TemBERTure_CLS_ results, we developed TemBERTure_Tm_, to predict protein melting temperature (Tm) from its primary sequence. Extracting the readily available protein melting temperature data from the Meltome Atlas, we again leveraged protBERT-BFD and adapter layers for training TemBERTure_Tm_. Even though the model achieved a seemingly high Pearson correlation of 0.78, a more detailed analysis revealed a clear limitation (Figure 4A). The predicted temperatures displayed a surprising bimodal distribution, concentrated around non-thermophilic (below 60°C) and thermophilic (above 80°C) ranges. This suggests a bias towards classifying temperatures into these broad categories rather than accurately predicting the melting points. This bias agrees with the weak correlation within each class (0.41 for non-thermophilic, −0.33 for thermophilic) and high accuracy (82%) of TemBERTure_Tm_ as a classifier using a 70°C threshold. Moreover, TemBERTure_Tm_ displayed significant variability among replicates trained with different random seeds, suggesting instability and limitations within the training process.

**Figure 4.**
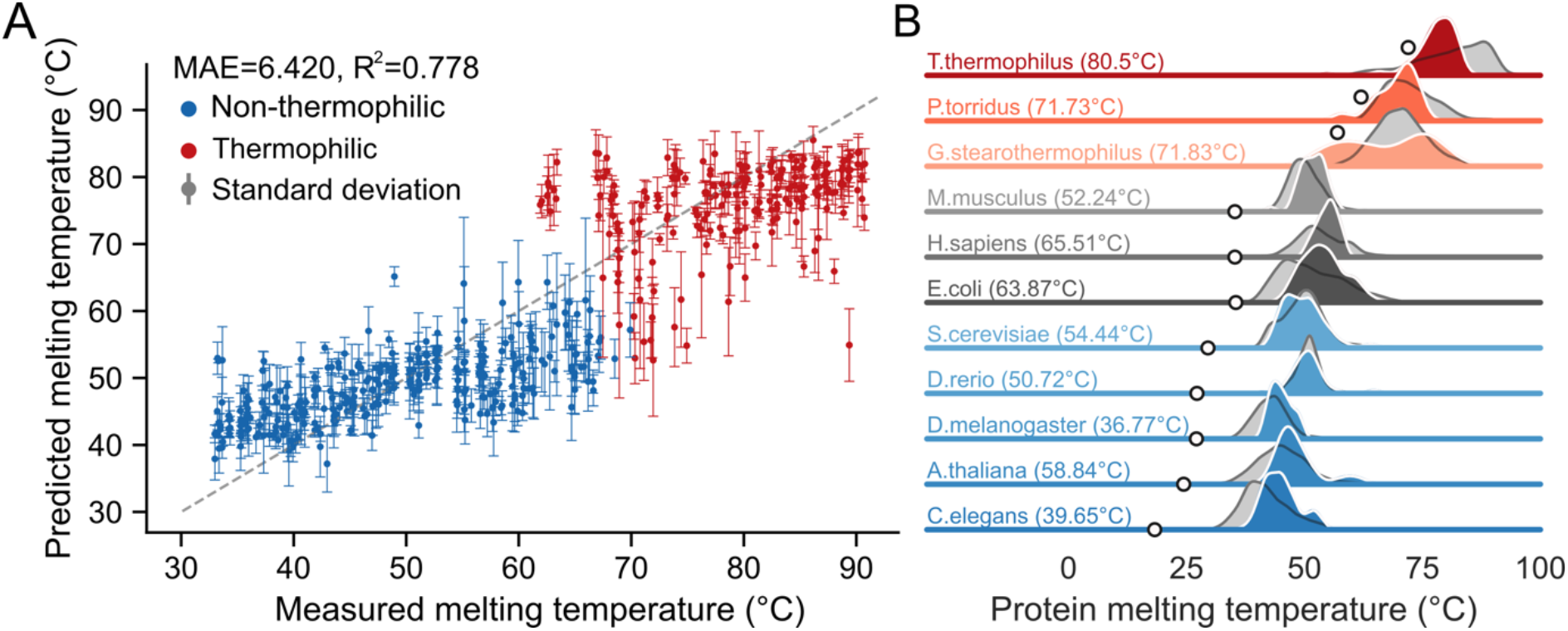
Predicted melting temperatures. A) Scatter plot comparing the measured melting temperatures to predicted melting temperature. Each point is colored base on the thermal category (blue: non-thermophilic and red: thermophilic). The dashed gray line represents a perfect prediction. Standard deviations are calculated from the predictions of three replicates. B) Distributions of melting temperature for various organisms, represented by a colored gradient ranging from red (high growth temperature) to blue (low growth temperature). The measured melting temperature distributions are shown in gray, while the predicted distributions using TemBERTure_TM_ are shown in color. Gray circles mark the growth temperatures of each organisms and the temperatures noted in parentheses indicating the average melting temperatures of the organism’s proteome.

Given the limited size (around 30,000 sequences) of the Metabolome Atlas dataset, we explored transfer learning. We hypothesized that pre-trained adapter weights from TemBERTure_CLS_, which captured thermal class features, could improve TemBERTure_Tm_ regression performance. Our approach involved replacing the random initialization of the adapter layers with weights from various stages of the classification training process. Since TemBERTure_Tm_ prediction followed a bimodal distribution, we chose different training stages for the adapter weights, aiming to balance leveraging learned thermal features and enabling the regression to move beyond this bias. However, this approach did not yield any significant improvements in performance.

In order to improve the performance, we explored diverse ensembling strategies (see Extended methods in Supplementary material). First, we established an upper bound on achievable performance using an oracle approach. From all TemBERTure_Tm_ variations, the oracle selected the prediction from all TemBERTure_Tm_ variations that was closest to the experimentally measured melting temperature. This yielded a best-case scenario with a MAE of 2.64°C and an R^2^ of 0.94 on the test set, highlighting the potential of the underlying approach. However, the ensemble techniques only led to marginal changes in performances (Table S7). A more promising approach involved leveraging thermal class information. We first predicted a protein’s class (non-thermophilic or thermophilic) using TemBERTure_CLS_ to predict the thermal class (non-thermophilic or thermophilic) of the protein sequence. Then, we selected a subset of best performing TemBERTure_Tm_ models for each class. This resulted in a combination of 5 models for non-thermophilic predictions (all transfer learning) and 2 models for thermophilic predictions (Table S7), i.e., one with random weights and one with partial first-epoch weights. This highlights the importance of incorporating class information, achieving a decrease in MAE (6.31°C) and an increase in R^2^ (0.78) on the test set compared to other ensembling techniques.

Despite limitations in predicting individual melting point prediction, TemBERTure_Tm_ showed promise in capturing broader thermal properties. We used the model to predict melting temperatures for unmeasured proteins from organisms within the Metabolome Atlas. Interestingly, the predicted distribution mirrored the known distribution of measured melting temperatures across diverse organisms (Figure 4B). This suggests that, although TemBERTure_Tm_ has some difficulties in predicting individual values, it still might capture underlying patterns related to protein thermostability across species.

### 2.4 Interpretability

To explore the intricate relationships between amino acid properties and thermostability, we conducted an analysis of the attention mechanisms in the TemBERTure_CLS_ model. Attention mechanisms offer an interpretable scoring function, highlighting segments of the input sequence that are most important for a particular prediction by assigning them higher scores. In the context of TemBERTure_CLS_, this would allow for a comprehensive identification of crucial amino acids and regions within a sequence that may influence the thermostability prediction. We defined High-Attention Score (HAS) regions as exceeding the interquartile range (IQR) of attention values across the entire sequence. All analyses were performed using the first replica of TemBERTure_CLS_.

#### Effect of fine-tuning

To investigate the impact of fine-tuning on the model’s attention patterns, we compared the frequencies of HAS amino acids between the pre-trained protBERT-BFD model and TemBERTure_CLS_. We hypothesized that changes in HAS frequencies might correlate with features linked to thermostability. Although the overall attention scores remained remarkably similar between the two models, we observed a shift in the frequency of HAS for specific amino acids (Figure 5A). For thermophilic proteins, leucine, arginine, and alanine appeared more frequently as HAS, whereas the frequency only increased for leucine in non-thermophilic sequences (Figure S1).

**Figure 5.**
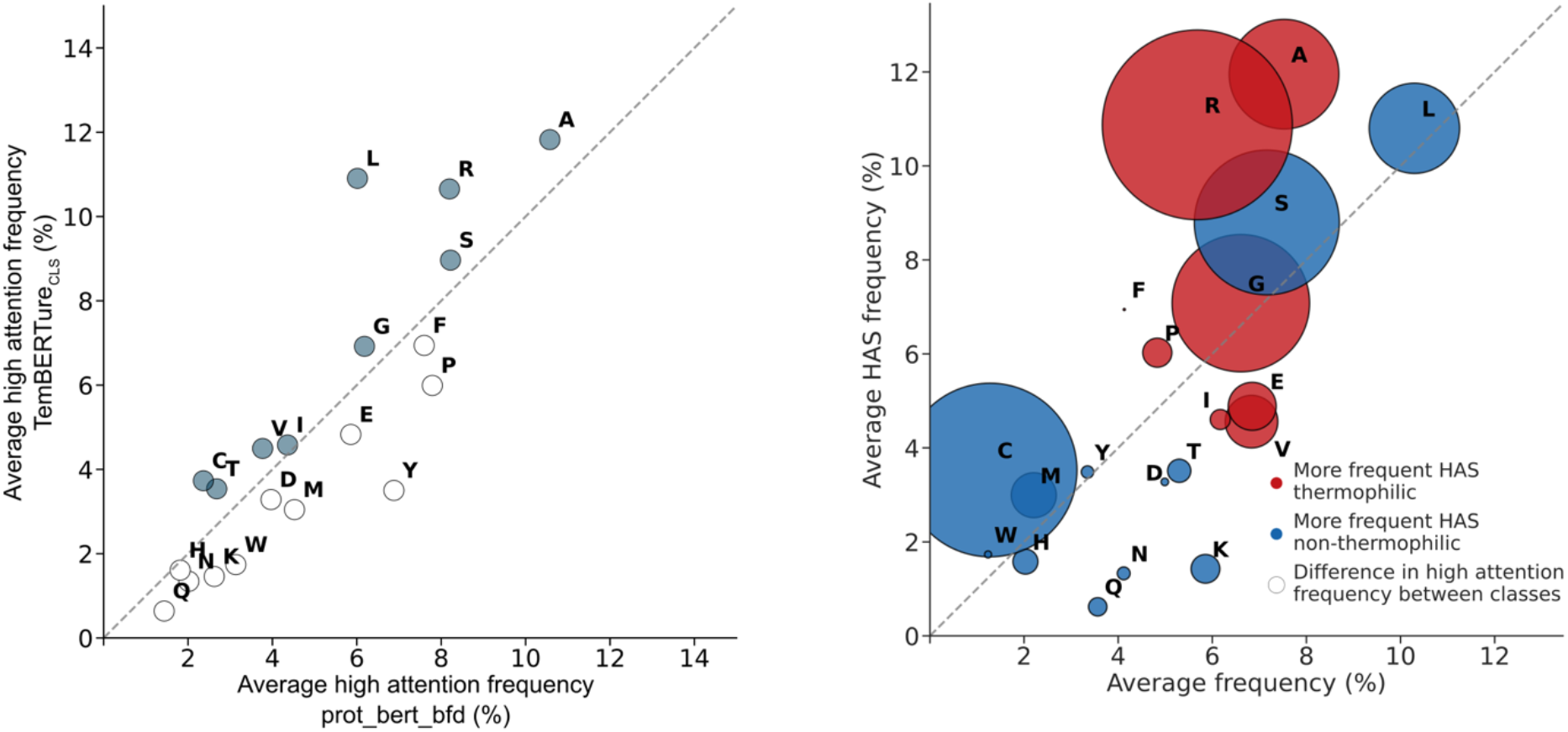
Frequency of high attention score (HAS) by amino acid. A) Scatter plot comparing the frequency of HAS amino acids of the pre-trained ProtBERT-BFD model to TemBERTure_CLS_. Each point represents an amino acid and is colored in gray if the frequency of HAS increased in TemBERTure _CLS_. B) Bubble plot comparing the frequency of each amino acid in the test set to its HAS frequency. Red bubble indicate that the frequency of HAS is higher for thermophilic and blue bubbles for non-thermophilic. Each bubble is scaled to the difference in frequency between both classes.

#### Amino acids enrichment

We conducted a more in-depth analysis by comparing the enrichment levels of each amino acid within the protein sequences with their natural occurrence frequencies. We calculated the background frequency of each amino acid in the TemBERTure_DB_ test set and compared it to the frequency at which they appeared as HAS (Figures 5B and S2). This analysis revealed distinct patterns between thermophilic and non-thermophilic proteins. For example, we observed an increase in HAS frequency for several hydrophobic residues, such as alanine, phenylalanine and leucine, which potentially reflect their role in stabilizing the protein core through tight packing. Interestingly, cysteine, which is known for forming stabilizing disulfide bridges and coordinating metals^46^, received higher attention in non-thermophiles. Glutamine and Asparagine, susceptible to deamidation at high temperatures^47–49^, showed decreased HAS, in agreement with their expected scarcity in these organisms. TemBERTure_CLS_ also showed a clear preference for different charged amino acids, with an increase in HAS for arginine and a decrease in HAS for lysine. However, it is crucial to underscore the potential complexity in interpreting HAS scores. An increase in high-attention scores (HAS) might suggest functional importance; however, their interpretation requires caution due to dependence on the local amino acid environment. Conversely, decreased HAS for specific amino acids might not indicate a negative impact, but rather reflect the model’s focus on their specific critical interactions within the protein structure.

#### Structural analysis

In order to gain some structural insights from the attention scores, we analyzed 17 pairs of homologous thermophilic and non-thermophilic proteins correctly classified by TemBERTure_CLS_. These pairs shared moderate sequence similarity (identity score: 0.28 - 0.54). Although the overall attention patterns between homologous proteins showed some correlation, the HAS amino acids exhibited more variability. Between homologous proteins, the model assigned a similar number of HAS to both conserved and non-conserved amino acids (Figures 6A and S3). Interestingly, the specific amino acids receiving HAS often differed between homologs, even in conserved regions. This is further supported by the presence of many HAS within insertion regions, highlighting the model’s ability to focus on regions beyond the conserved core for thermostability prediction.

**Figure 6.**
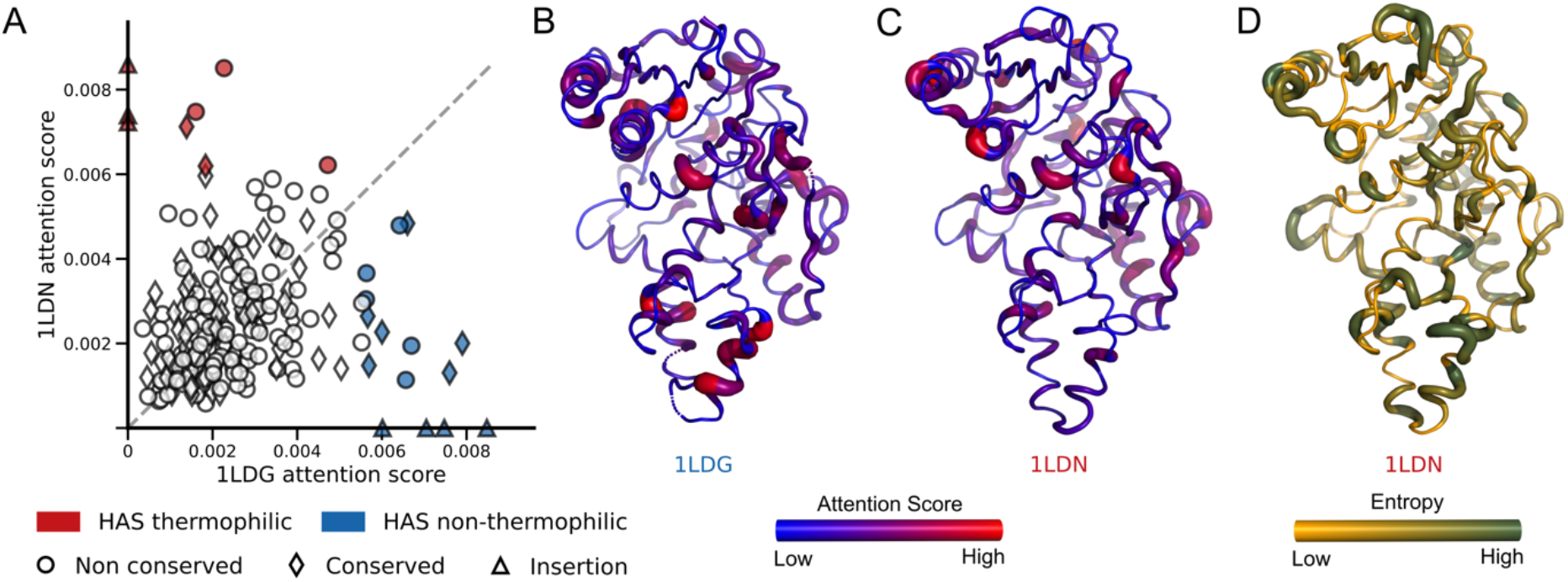
Representative structural analysis of attention scores. (A) Scatter plot comparing the attention scores assigned by the TemBERTure_CLS_ model to individual amino acids in two homologous protein structures (PDB ID: 1LDN [thermophilic] and 1LDG [non-thermophilic]) with 46% sequence identity. . Each marker represents an amino acid, categorized by its conservation level: circles for non-conserved, diamonds for conserved, and triangles for insertions. HAS amino acids in the thermophilic structure are highlighted in red, while those in the non-thermophilic counterpart are highlighted in blue. (B) and (C) Cartoon representation of both protein structures. The width and color indicate the attention score values, with regions with higher attention scores appearing thicker and redder. D Cartoon representation of 1LDN colored based on the entropy at each amino acid position position. Higher entropy (green, thicker regions) indicates greater sequence variability.

To understand how TemBERTure_CLS_ leverages structural information beyond sequence similarity, we mapped the attention scores directly onto protein structures (Figure 6B, C, and S4). Higher attention scores localized similarly across homologs, regardless of sequence entropy (Figure 6D). Notably, higher attention scores often resided in helical regions and the protein core, potentially revealing the prioritization of structurally important elements for predicting thermostability.

## 3. Discussion

Protein thermostability is crucial for various applications in biotechnology and biology. Traditional experimental methods for assessing it are laborious, expensive, and prone to errors. Here, we developed a new set of tools which allowed us to explore the potential of Deep Learning models to predict protein thermostability. Our study highlights the critical role of data diversity in training robust models. We observed significant performance improvement with datasets encompassing a wider range of sequences from various organisms. Conversely, insufficient diversity, as seen in the BacDive derived dataset, led to models that struggled with challenging test sets. This emphasizes the need for a holistic approach to data curation, in order to ensure balanced representation of diverse species in the training data.

Although the Meltome Atlas presents an impressive number of melting temperatures, it suffers from certain biases, in particular, the data primarily represents non-thermophilic organisms with a temperature gap between 60 – 70°C. Interestingly, TemBERTure_TM_’s predictions, while not accurate for absolute melting temperatures, captured the overall distribution of melting temperatures observed across different species in the dataset. This suggests the model might have prioritized recognizing the species origin of the sequence rather than intrinsic thermostability features. This agrees with previous findings showing that sequence embeddings from language models can already capture these broad differences between thermophilic and non-thermophilic organisms^38^. Additionally, the presence of thermostable proteins within non-thermophilic proteomes further underscores the limitations of using growth temperature alone as a thermostability proxy.

Various statistical approaches have attempted to identify important changes in amino acid composition linked to thermostability^13,22,50–54^. However, such analyses heavily depend on dataset curation, leading to contradictory results. Furthermore, while certain biophysical properties of residues may elucidate their prevalence in thermostable proteins, thermophilicity is a multifaceted attribute influenced by the positioning and microenvironment of amino acids within the protein. This study presents the concept of leveraging attention scores to gain more nuanced insights into protein thermostability. Even though we observed some global trends consistent with previous analyses (e.g., enrichment of specific amino acids), TemBERTure_CLS_ also highlighted the value of analyzing these interactions within the context of the 3D protein structure. However, our findings suggest that the present attention scores still need to be refined, since they capture both thermostability-related features and organism-specific characteristics. Further research is needed to refine them for a more precise understanding of protein thermostability.

In conclusion, this work sheds light on the limitations of current approaches for predicting protein thermostability. It introduced new avenues for exploration. which highlighted the importance of using diverse training data, extending the analysis beyond single-species, and exploiting important features of the models, such as attention scores. Based on our results, future research can develop even more robust and informative methods for predicting protein thermostability.

## 4. Materials and Methods

This section is composed of four main parts. Part 1 outlines the workflow for establishing comprehensive curated databases of thermophilic and non-thermophilic protein sequences sourced from various experiments and data collection, with TemBERTure_DB_ as the primary training resource and two additional databases used for bias and generalization assessment. The second and third subsection describes the architecture and training of TemBERTure_CLS_ and Temberture_Tm_. The last subsection provides the technical details used for the analyses.

### 4.1 Database creation

#### a. TemBERTure_DB_

TemBERTure_DB_ leveraged data from the Meltome Atlas experiment^10^. We obtained pre-processed protein sequences from the ProtStab2 dataset^33^. These sequences were supplemented by retrieving all sequences from UniProtKB^43^ corresponding to the same thirteen organisms as in the Meltome Altas. To address the class imbalance between thermophilic and non-thermophilic sequences, we enriched the thermophilic dataset by sourcing additional data from the BacDive database^44^. Here, we classified sequences based on the growth temperature of their respective organisms: thermophilic (>60°C) and non-thermophilic (<30°C). Protein sequences were retrieved for each organism from the NCBI database^55^. Ambiguous and short (< 30 amino acids) sequences were excluded. MMseqs was then employed to cluster the sequences within each dataset, using a threshold of 50% for thermophilic and 80% for non-thermophilic. To further address the class imbalance, we augmented the non-thermophilic dataset with challenging examples. These examples were retrieved from non-thermophilic organisms (BacDive) and exhibited high sequence similarity (80% < identity < 95%) to the thermophilic sequences. The final TemBERTure_DB_ was stored as an SQL database facilitating efficient data retrieval for downstream analyses (Table S1).

#### b. BacDive

Within the BacDive database, organisms were classified based on growth temperature: thermophilic (>60°C) and non-thermophilic (<30°C). Protein sequences were then retrieved for each organism from the NCBI database, and ambiguous or short sequences (<30 amino acids) were excluded. Given the substantial disparity between the number of non-thermophilic and thermophilic sequences, we used MMseqs in cascading mode to cluster the non-thermophilic sequences. We then undersampled the centroids (representatives of each cluster) to align with the number of thermophilic centroids identified using MMseqs with a 50% identity threshold (Table S5).

#### c. Meltome

We leveraged data curated within TemBERTure_DB_ and excluded the non-thermophilic counterparts of the high-similarity sequence pairs retrieved from the BacDive database (Table S6).

#### Splitting

For model training, we partitioned the datasets into an 80:10:10 ratio for the training, validation, and test sets, respectively. To mitigate any potential information leakage between sets, all sequences were clustered with MMseqs at a 50% identity threshold. Centroids and their corresponding clusters were then assigned to the same split.

For the regression task, we exclusively used the initial Meltome dataset. Melting temperatures were categorized into temperature bins of 10°C, and 10 data points from each temperature bin were randomly selected for both the test and validation sets. To address the imbalance in the distribution of melting temperatures within the training set, we implemented a combination of undersampling and oversampling techniques. Temperature bins with an abundance of data points (40 – 55 °C) were undersampled, whereas bins with a scarcity of data points (20 – 40°C and 60 – 90°C) were oversampled. This approach ensured a balanced number of data points across all temperature bins.

### 4.2 TemBERTure_CLS_

TemBERTure_CLS_ (Figure 1B) is a sequence-based classifier that takes the amino acid sequence as input and outputs the corresponding thermal class of the protein along with its associated score. It was built on top of the pre-trained protBERT-BFD model^35^, a BERT model composed of 30 layers, 16 heads, and 1024 hidden layers and trained on over 2 billion protein sequences from the BFD100^56,57^ dataset. In order to reduce the number of trainable parameters and enhance the efficiency of the training process, we opted for an adapter-based fine tuning technique^41,42^, where light weight bottleneck layers are inserted between each transformer layer.

TemBERTure_CLS_ was thus implemented as a BertAdapterModel with Pfeiffer adapters^58^ configuration using the PyTorch framework via adapters^42^ library. It was initiated with the proBERT-BFD^35^ weights through the HuggingFace API^59^ and the Pfeiffer adapter architecture layers were added after the feed-forward block of each Transformer layer^60 61^. In this way we reduced the number of trainable parameters from 420 million to 5 million.

#### Training

Protein sequences were tokenized at the amino acid level utilizing the protBERT-BFD^35^ tokenizer, with all sequences truncated to a maximum length of 512. For each dataset, a separate hyperparameter search was carried out to optimize the training and architecture of the model (Table S8). This hyperparameter search was performed through the use of W&B Sweeps^62^ grid hyperparameter search. The adapter training was carried out for a maximum of 20 epochs for each dataset with a batch size of 16, using AdamW optimizer^63^ with default Hugging Face^59^ configuration. The model that achieved the lowest validation loss was then saved for evaluation. To ensure model robustness, the final configuration of each model was trained three times under identical conditions, varying only the random seed. This approach allowed us to assess the model’s independence from specific random seeds and to confirm its reliability across different runs. All models were trained on a single NVIDIA A100 80G GPU.

### 4.3 TemBERTure_Tm_

TemBERTure_Tm_ is a sequence-based regression model designed to predict the protein melting temperature (Tm) directly from its amino acid sequence. This model has the same underlying architecture configuration and tokenization as TemBERTure_CLS_, with a regression head. Leveraging the pre-trained protBERT-BFD model, we adopted again an adapter-based fine-tuning technique to reduce trainable parameters.

#### Training

The model was trained on a curated dataset created specifically for predicting protein melting temperatures, based on TemBERTure_DB_. All sequences are truncated to a maximum length of 512. The training was carried out for a maximum of 200 epochs for each run with a batch size of 16 and using AdamW optimizer ^63^ with default Hugging Face ^59^ values. We conducted, with W&B Sweeps ^62^, an extensive search to identify the optimal configuration of the regression head (Table S9). We then explored various weight initialization approaches for the model. In addition to random initialization, we investigated transfer learning from TemBERTure_CLS_ at different training stages. This involved introducing classifier weights at 25%, 50%, 75%, and 100% of the first epoch, along with weights from the fully trained classifier. To assess model stability and consistency across random initializations, all models were trained three times with different random seeds. For each configuration, the model achieving the lowest validation loss was saved for further evaluation. All training runs utilized a single NVIDIA A100 80G GPU.

### 4.4 Analyses

#### Ensemble Evaluation for Melting Temperature Prediction

To improve prediction accuracy, we evaluated different ensembles of models on the validation set. We built these ensembles by selecting subsets of the initial 18 models. These 18 models encompassed all distinct initialization methods (random and transfer learning with TemBERTure_CLS_ weights) and their replicates. We investigated three ensemble approaches: greedy algorithm, weighted ensemble, and a method leveraging TemBERTure_CLS_. Additionally, we experimented with various averaging techniques (standard deviation and interquartile range) to combine predictions and identify the optimal value for each data point. Overall, these ensemble strategies aimed to harness the strengths of multiple models and achieve a configuration effective across a broad temperature range. Detailed descriptions are provided in the Extended Methods in the supporting information.

#### High attention score

The interquartile range (IQR) method was used to identify amino acids within a protein sequence with a high attention score (HAS). We calculated a threshold by adding 1.5 times the IQR to the third quartile (Q3) of the attention scores. Attention scores exceeding this threshold are flagged as outliers, indicating a noticeably high attention score (HAS) and potentially significant influence on the model’s decisions.

