## Supplementary Materials for "TemBERTure: Advancing protein thermostability prediction with Deep Learning and attention mechanisms"

+ Authors contributed equally

ORCID

Chiara Rodella: 0009-0002-2127-6594

Symela Lazaridi: 0009-0003-3323-7215

Thomas Lemmin: 0000-0001-5705-4964

### Extended Methods

#### Ensemble Evaluation for Melting Temperature Prediction

To enhance a better melting temperature prediction of $TemBERTure_{TM}$, we evaluated model ensembles on the validation set. These ensembles were constructed by selecting a subset of the initial 18 models, which covered all distinct initialization methods (random and transfer learning with $TemBERTure_{CLS}$ weights) and their duplicates. We explored three ensemble approaches: greedy algorithm, weighted ensemble, and a method leveraging $TemBERTure_{CLS}$. Additionally, we experimented with various averaging techniques (standard deviation and interquartile range) to combine predictions and identify the optimal value for each data point. Overall, these ensemble strategies aimed to harness the strengths of multiple models and achieve effectiveness across a broad temperature range.

*Averaging*

The predictions were averaged across all replicas, resulting in an average melting temperature from 18 models per observation.

*IQR and standard deviation*

The predictions of all models were aggregated, We then identified and removed outliers using either the interquartile range (IQR) or a 3 standard deviation threshold. The remaining inliers are averaged for a final prediction.

*Greedy search ensemble*

The greedy ensemble approach aimed to identify an optimal combination of models to minimize Mean Absolute Error (MAE) on the validation set. We began by initializing the ensemble with the model having achieved the lowest MAE. We then iteratively evaluated the performance (based on MAE) of adding one additional model to the ensemble. If the average prediction resulted in a lower overall MAE, we updated the ensemble to include that model. The process continued until no further improvement for a maximum of 3 iterations. We also carried out a similar approach, setting the maximum number of models to 5.

*Classification-Based Ensemble*

This method leverages a two-stage ensemble approach for predicting melting temperature. Each sequence in the validation set is assigned a class label (thermophilic or non-thermophilic) using the $TemBERTure_{CLS}$ model. For each class, a greedy search ensemble approach was employed to select the set of models that minimized the Mean Absolute Error (MAE) on the corresponding target melting temperature values.

**Table S1: Summary of data processing pipeline for TemBERTure$_{DB}$.**

| Step | Description | Input | Output | Source of Data |
|---|---|---|---|---|
| 1. Sequence Acquisition | Retrieve sequences for organisms in Meltome Atlas [10] | 13 Organisms | 4,283,681 sequences (40,970 thermophilic, 4,242,711 non-thermophilic) | UniProtKB [1] |
| 2. Initial Centroid Generation | Cluster sequences using MMseqs (>50% identity) | Sequences from Step 1 | 574,600 non-thermophilic, 13,330 thermophilic centroids | Sequences from Step 1 |
| 3. Data Balancing (Thermophilic) | Enrich thermophilic data | Organisms with growth temperature >60°C | 467 thermophilic organisms (bacteria & archaea) | BacDive database [2] |
| 4. Protein Sequence Retrieval (Thermophilic) | Retrieve protein sequences for organisms and filter out short sequences (<30 aa) | Thermophilic organisms (Step 3) | 886,269 thermophilic protein sequences | NCBI database [3] |
| 5. Centroid Generation (Thermophilic) | Cluster thermophilic sequences (MMseqs, >50% identity) | Thermophilic sequences (step 4) | 272,658 thermophilic | Thermophilic sequences from step 4 |
| 6. Data Balancing (Non-thermophilic) | Enrich non-thermophilic data | Organisms with growth temperature <30°C | 11,028 non-thermophilic organisms (10,988 bacteria and 40 archaea) | BacDive database [2] |
| 7. Protein Sequence Retrieval (Non-Thermophilic) | Retrieve protein sequences for organisms and filter out short sequences (<30 aa) | Non-thermophilic organisms (Step 6) | 35,247,992 protein sequences for non-thermophilic organisms | NCBI database [3] |
| 8. Centroid Generation (Non-thermophilic) | Cluster non-thermophilic sequences (MMseqs, >80% identity) | Non-thermophilic sequences (Step 7) | 15,756,711 non-thermophilic centroids | Non-thermophilic sequences from step 6 |
| 9. Challenging Pairs (Non-thermophilic) | Derive only non-thermophilic sequences that are similar at 80-95% identity to thermophilic sequences | Thermophilic and non-thermophilic sequences (steps 5 & 8) | 37,203 challenging sequences | Thermophilic and non-thermophilic sequences from steps 5 and 8 |
| 10. TemBERTure$_{DB}$ | Merge and balance data from Meltome Atlas, Bacdive, and UniProt databases derived from previous steps | All previous data | 241,472 thermophilic, 250,605 non-thermophilic, | All previous data |

**Table S2: Number of sequences in the TemBERTure$_{DB}$ classifier and regression datasets**

| | Classifier | | | Regression | | |
|---|---|---|---|---|---|---|
| | **Train** | **Validation** | **Test** | **Train** | **Validation** | **Test** |
| **Thermophilic** | 241472 | 26649 | 26643 | 10296 | 203 | 209 |
| **Non-thermophilic** | 250605 | 22404 | 22380 | 19780 | 319 | 322 |

**Table S3: Comparison between our TemBERTure$_{CLS}$ model and other state-of-the-art models on the TemBERTure$_{DB}$ test set.**

| Models | Thermophilic Recall | Non-thermophilic Recall | Average Accuracy | F1-Score | Non-thermophilic Precision | Thermophilic Precision | Matthews correlation coefficient |
|---|---|---|---|---|---|---|---|
| **TemBERTure$_{CLS}$** | 0.889 | 0.897 | 0.893 | 0.900 | 0.872 | 0.911 | 0.784 |
| **iThermo** [4] | 0.633 | 0.812 | 0.723 | 0.707 | 0.650 | 0.800 | 0.448 |
| **SCMTPP** [5] | 0.477 | 0.847 | 0.662 | 0.594 | 0.577 | 0.788 | 0.344 |
| **ThermoPred** [6] | 0.636 | 0.798 | 0.717 | 0.704 | 0.648 | 0.789 | 0.435 |
| **BertThermo** [7] | 0.700 | 0.809 | 0.755 | 0.753 | 0.694 | 0.814 | 0.509 |
| **TemStaPro$_{-minor-30}$** [8] | 0.661 | 0.968 | 0.786 | 0.786 | 0.662 | 0.968 | 0.629 |

**Table S4: TemBERTure$_{CLS}$ generalization ability**. TemBERTure$_{CLS}$ performance on available test sets, excluding sequences with over 50% identity our TemBERTure$_{DB}$ validation and training sets. To account for the large class imbalance in the filtered dataset, we use macro-averaging for F1-score, recall, and precision.

| Models | Non-overlapping Thermophilic Sequences | Non-overlapping Non-thermophilic Sequences | F1-score | Recall | Precision[4] |
|---|---|---|---|---|---|
| **iThermo** | 65 | 505 | 0.773 | 0.924 | 0.729[8] |
| **TemStaPro$_{-minor-30}$** | 1495 | 10849 | 0.738 | 0.891 | 0.705 |

**Table S5: Summary of data processing pipeline for $BacDive_{DB}$.**

| Step | Description | Input | Output | Source of Data |
|---|---|---|---|---|
| 1. Protein Sequence Retrieval (Thermophilic) | Retrieve protein sequences for organisms with growth temperature >60°C and filter out short sequences (<30 aa) | Organisms with growth temperature >60°C | 900,859 thermophilic protein sequences | BacDive database [2]<br>NCBI database [3] |
| 2. Centroid Generation (Thermophilic) | Cluster thermophilic sequences (MMseqs, >50% identity) | Thermophilic sequences (step 2) | 278,894 thermophilic | Thermophilic sequences from step 1 |
| 3. Protein Sequence Retrieval (Non-Thermophilic) | Retrieve protein sequences for organisms with growth temperature <30°C and filter out short sequences (<30 aa) | Organisms with growth temperature <30°C | 40,032,534 protein sequences for non-thermophilic organisms | BacDive database [2]<br>NCBI database [3] |
| 4. Reduce the number of Non-thermophilic sequences | Cluster non-thermophilic sequences (MMseqs cascading clustering) | Non-thermophilic sequences (step 3) | 3,985,266 non-thermophilic centroids and randomly select the same amount as the thermophilic dataset (900,859 non-thermophilic) | Non-thermophilic sequences from step 3 |
| 5. Centroid Generation (Non-thermophilic) | Cluster non-thermophilic sequences (MMseqs, >50% identity) | Non-thermophilic sequences (step 4) | 861,960 non-thermophilic | Non-thermophilic sequences from step 4 |
| 6. $BacDive_{DB}$ | Merge and balance data from previous steps | All previous data | 278,894 thermophilic, 278,894 non-thermophilic, | All previous data |

**Table S6: Summary of data processing pipeline for Meltome$_{DB}$.**

| Step | Description | Input | Output | Source of Data |
|---|---|---|---|---|
| 1. Sequence Acquisition[9] | Retrieve sequences for organisms in Meltome Atlas | 13 Organisms | 4,283,681 sequences (40,970 thermophilic, 4,242,711 non-thermophilic) | UniProtKB [1] |
| 2. Initial Centroid Generation | Cluster sequences using MMseqs (>50% identity) | Sequences from Step 1 | 574,600 non-thermophilic, 13,330 thermophilic centroids | Sequences from Step 1 |
| 3. Data Balancing (Thermophilic) | Retrieve protein sequences for organisms with growth temperature >60°C and filter out short sequences (<30 aa) | Organisms with growth temperature >60°C | 881,831 thermophilic protein sequences | BacDive database [2]<br>NCBI database |
| 4. Centroid Generation (Thermophilic) | Cluster thermophilic sequences (MMSeqs, >50% identity) | Thermophilic sequences (step 4) | 286,123 thermophilic | Thermophilic sequences from step 4 |
| 5. Meltome$_{DB}$ | Merge and balance data from Meltome Atlas, Bacdive, and UniProt databases derived from previous steps | All previous data | 286,123 thermophilic, 316,929 non-thermophilic, | All previous data |

**Table S7: Ensemble performances on the regression validation set**. Mean Absolute Error (MAE) computes the discrepancy between the predicted melting temperatures and the actual observed values. The coefficient of determination, denoted as R2, assesses the goodness of fit of the model in capturing the variability of the melting temperature.

| | | All | Non-thermophilic | Thermophilic |
|---|---|---|---|---|
| Oracle | MAE | 3.369 | 3.53 | 3.117 |
| | R2 | 0.91 | 0.715 | 0.595 |
| Greedy algorithm (11 models) | MAE | 7.004 | 6.981 | 7.041 |
| | R2 | 0.748 | 0.25 | -0.274 |
| Weighted ensemble (5 models) | MAE | 7.065 | 7.028 | 7.124 |
| | R2 | 0.741 | 0.24 | -0.343 |
| Classification-based - non-thermophilic (5 models)- thermophilic (5 models) | MAE | 6.769 | 6.834 | 6.668 |
| | R2 | 0.761 | 0.261 | -0.135 |
| Average after IQR outliers identification | MAE | 7.042 | 7.01 | 7.092 |
| | R2 | 0.746 | 0.247 | -0.297 |
| Average after std outliers identification | MAE | 7.057 | 7.027 | 7.104 |
| | R2 | 0.746 | 0.248 | -0.297 |
| Average within best replicas | MAE | 5.182 | 5.237 | 5.096 |
| | R2 | 0.837 | 0.505 | 0.205 |
| Average within all the replicas | MAE | 7.056 | 7.023 | 7.107 |
| | R2 | 0.746 | 0.249 | -0.297 |

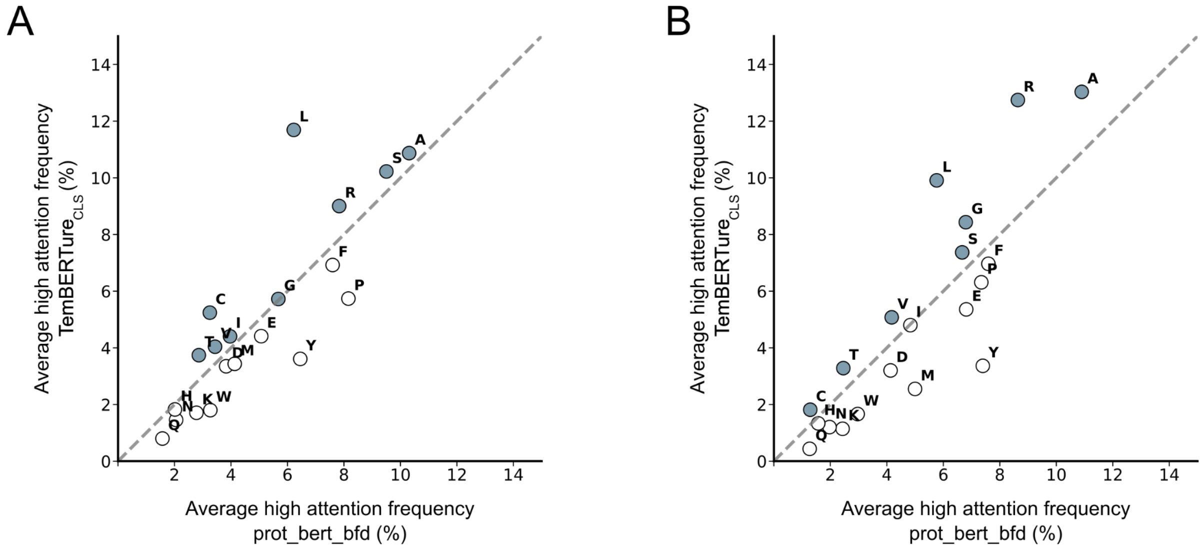


**Figure S1: Effect of fine-tuning on amino acid HAS frequency.** Scatterplot comparing the high attention frequency of amino acids as identified by the pretrained protBERT-BFD [10] model (x-axis) versus the fine-tuned TemBERTure$_{CLS}$ model (y-axis) respectively for non-thermophilic sequences (A) and thermophilic sequences (B). Each point represents an amino acid, with its position reflecting the change in attention frequency after fine-tuning. Amino acids located above the diagonal have gained attention in the fine-tuned model and are colored in gray.

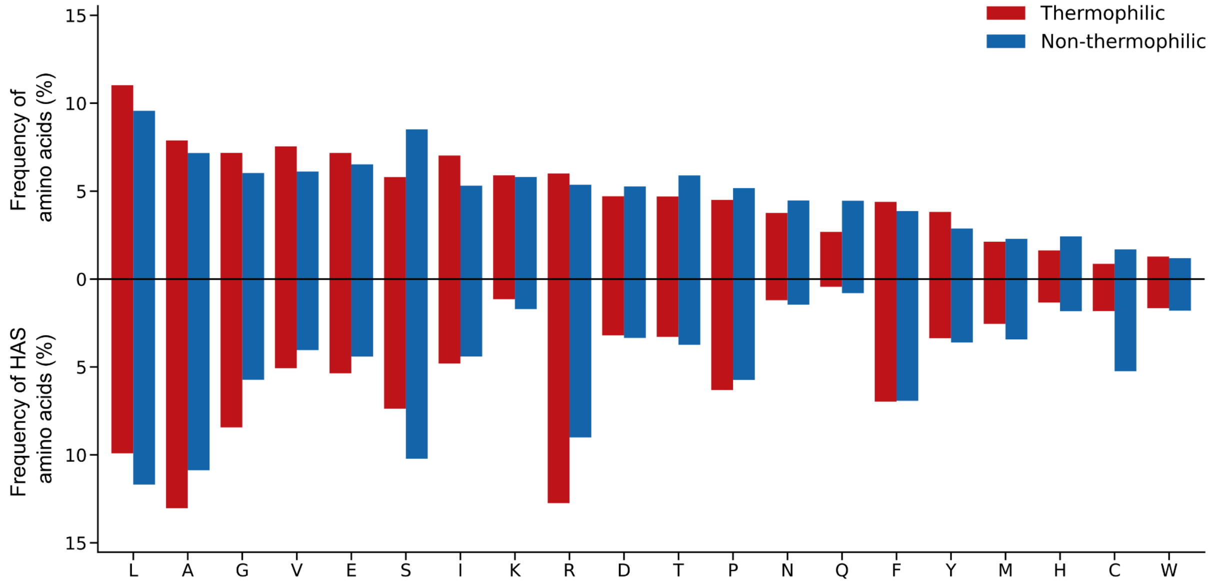


**Figure S2. Amino acid frequency and HAS frequency.** The bar chart presents a dual-layered comparison: the upper segment displays the frequency of individual amino acids within the TemBERTure$_{DB}$ test set, while the lower segment focuses specifically on the frequency of HAS amino acids. Red bars represent the prevalence of amino acids in thermophilic proteins, while blue bars denote their occurrence in non-thermophilic proteins.

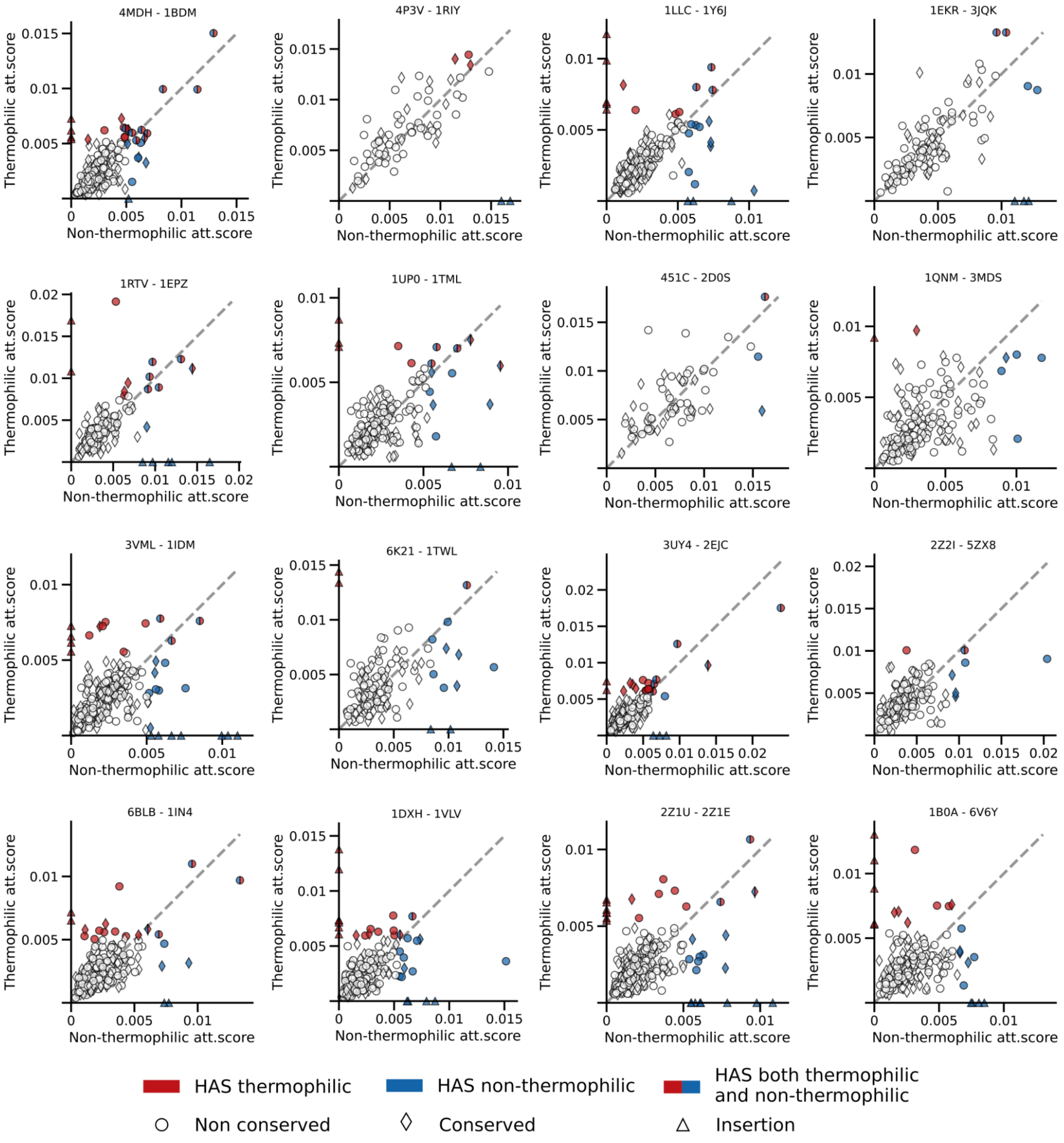


**Figure S3: Comparison of TemBERTure$_{CLS}$ attention scores on pairs of protein homologs**.

Scatter plots illustrate the attention scores between thermophilic and non-thermophilic paired PDB structures. Each plot corresponds to a unique pair, denoted by their respective PDB IDs. Red and blue markers indicate HAS for thermophilic and non-thermophilic respectively, and diamonds and circles differentiate between conserved and non-conserved amino acids, and triangles represent insertions.

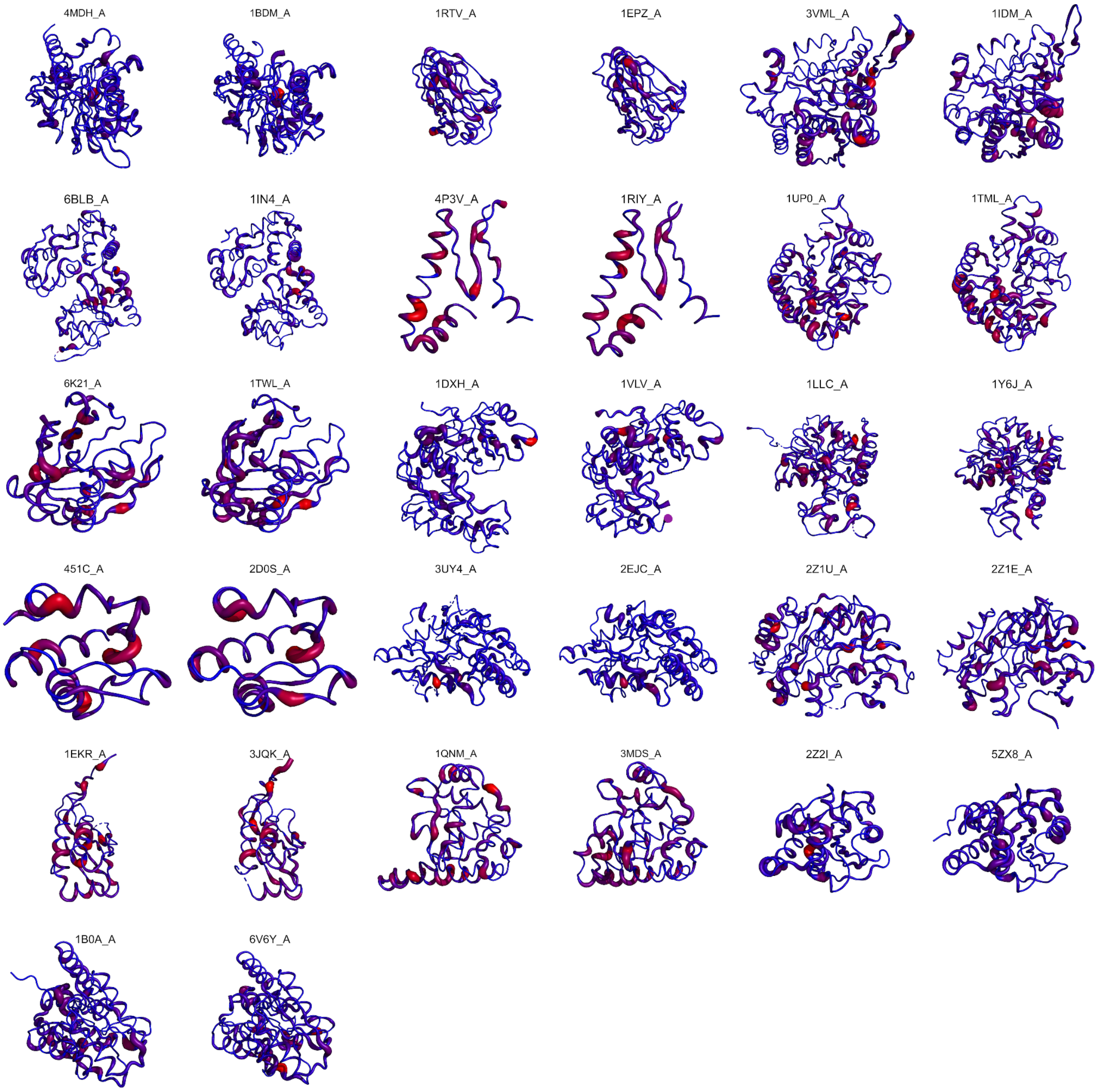


**Figure S4: Mapping of TemBERTure$_{CLS}$ attention score on protein structures.** Set of 16 pairs of homologous non-thermophilic (on the left of the pair) and thermophilic (on the right) protein structures extracted from the Protein Data Bank. Each pair of protein structures is depicted side by side for comparative analysis. Regions with a higher attention score are thickened and colored in red

**TableS8: TemBERTure$_{CLS}$ models hyperparameter tuning for different datasets.** The optimal setting is highlighted in bold

| Hyperparameters | Dataset | | |
|---|---|---|---|
| | BacDive | Meltome | TemBERTure$_{DB}$ (3 replicas) |
| Learning rate | 5e-3,5e-4,5e-5,**5e-6** | 5e-3,5e-4,5e-5,**5e-6** | 1e-3,1e-4,**1e-5**,1e-6, |
| Head dropout probability | **0**,0.1 | **0**,0.1 | 0,**0.1**,0.2,0.3,0.4 |
| Head activation function | **tanh** | **tanh** | **tanh** |
| Warmup ratio | **0** | **0** | **0**, 0.10 |
| Weight decay | **0** | **0** | **0**,0.10 |
| Lowest validation loss | **Epoch 9** | **Epoch 10** | **Epoch (4,4,5)** |
| Training time | **23 hours** | **32 hours** | **37 hours** |

**Table S9: TemBERTure$_{Tm}$ hyperparameter tuning**. Optimal parameters are shown in bold.

| | |
|---|---|
| Learning rate | 1e-2 **1e-3** 1e-4 1e-5 1e-2 |
| Head layers | 1,**2** |
| Head dropout probability | 0,0.1, 0.2, **0.3**, 0.4 |
| Head activation function | tanh, **ReLU**, LeakyReLU |
| Warmup ratio | **0**, 0.10 |
| Weight decay | **0**, 0.10 |
